# HtrA1 alters endothelial tube formation characteristics in an *in vitro* model

**DOI:** 10.1101/539304

**Authors:** Harmeet Singh, Guiying Nie

**Affiliations:** Implantation and Placental Developmental Laboratory, Centre for Reproductive Health, Hudson Institute of Medical Research, Clayton, Victoria 3168, Australia; Department of Molecular and Translational Sciences, Monash University, Clayton, Victoria 3800, Australia; Department of Biochemistry and Molecular Biology, Monash University, Clayton, Victoria 3800, Australia

**Keywords:** HtrA1, age-related macular degeneration, AMD, preeclampsia, endothelial dysfunction

## Abstract

High temperature requirement factor A1 (HtrA1) is a serine protease of the mammalian HtrA family. It is ubiquitously expressed with high levels in the placenta. Dysregulation of HtrA1 has been linked to a number of diseases, in particular age-related macular degeneration (AMD) and preeclampsia (PE) in which HtrA1 is significantly increased. AMD is the leading cause of irreversible visual impairment in older people, affecting millions across the globe. PE is a life-threatening pregnancy complication, affecting 2-7% of pregnant women worldwide. Although AMD and PE are very different diseases, both are associated with endothelial dysfunction and dysregulation of angiogenesis. Given HtrA1 is up-regulated in both AMD and PE, in this study we examined the impact of excessive HtrA1 on capillary tube formation of HUVECs as an *in vitro* angiogenesis model. HtrA1 at high concentrations significantly increased the total number of tube branch points and inter-tubular loops, but considerably decreased the mean tube length, resulting in more but much smaller tubes. However, these smaller tubes were incomplete/broken. These data demonstrated that high concentrations of HtrA1 altered endothelial tube formation characteristics of HUVEVs. Our results suggest that HtrA1 over-expression in AMD and PE may directly contribute to the endothelial dysfunction in these diseases.

## 1. Introduction

High-temperature requirement factor A1 (HtrA1) is a multi-domain protease of the mammalian HtrA family, which contains a well-conserved trypsin-like serine protease domain and a PDZ domain [1]. HtrA1 is expressed ubiquitously, but the highest level is found in the placenta [2, 3]. To date, dysregulation of HtrA1 has been implicated in a number of diseases such as cancer [4, 5], arthritis [6], age-related macular degeneration (AMD) [7], neurodegenerative and neuromuscular disorders [8], and pregnancy complication preeclampsia [9, 10]. Intriguingly, while HtrA1 is down-regulated in a number of cancers [4, 5, 11, 12], it is up-regulated in AMD [7, 13, 14] and preeclampsia (PE) [9, 10].

AMD is the leading cause of irreversible visual impairment in older people, affecting millions worldwide. AMD occurs in “dry” and “wet” forms [7, 13, 15]. Although dry AMD is more common, it does not typically result in blindness. In contrast, wet AMD, accounting for approximately 10% of AMD patients, can cause rapid vision loss and blindness [7, 13]. In wet AMD, new blood vessels form and break beneath the retina, and this leakage of blood causes permanent damage to surrounding tissue [7, 13].

Although AMD is a multifactorial disease, genome-wide association studies have linked a mutation in the HtrA1 gene with wet AMD [7, 13], which has been subsequently proven in many different populations [14, 16, 17]. It is further established that this mutation in HtrA1 enhances its expression and increases blood vessel branching [7]. Importantly, it is demonstrated that increased HtrA1 may be sufficient to induce wet AMD in mice [18].

PE is a multifactorial disorder of pregnancy that affects 3%-7% of pregnant women worldwide; it is one of the leading causes of maternal and neonatal mortality and morbidity [19]. PE is characterized by new onset hypertension, with proteinuria and/or maternal organ dysfunction after 20 weeks of gestation [20]. Based on the gestation age of symptom presentation, PE can be classified into early-onset (<34 gestation weeks) and late-onset (>34 weeks) subtypes [21]; the two differ in pathophysiology and disease severity, and in general early-onset PE poses more significant maternal risks with more severe perinatal outcomes compared to late-onset PE [21–23].

PE is associated with systemic endothelial dysfunction and generalised maternal inflammation, which are believed to give rise to the multi-systemic clinical symptoms [22, 24, 25]. Although the causes of PE are not completely understood, the placenta is known to play a key role. During pregnancy, the placenta releases many factors into the maternal circulation to modulate the maternal vasculature. In PE, placental production and secretion is altered, resulting in altered levels of pro- and anti-angiogenic factors and pro-inflammatory cytokines circulating in the maternal blood [24, 25], these abnormal levels of circulating factors are shown to play an essential role in inducing endothelial dysfunction in PE [26–29]. PE is thus considered as an endothelial disease.

HtrA1 is highly expressed in the placenta and closely associated with placental development [2, 3, 30]. Placental HtrA1 expression is significantly increased in PE, especially in the early-onset PE subtype [9, 31–33]. We have further reported that placental HtrA1 protein is secreted into the maternal circulation, and that the circulating levels of HtrA1 are also significantly elevated in early-onset PE compared to gestation-aged-matched controls [10].

Although wet AMD and early-onset PE are two very different diseases, both are characterized by dysregulation of angiogenesis and endothelial dysfunction, and in both diseases HtrA1 production is elevated. However, how HtrA1 affects the endothelial cells is not well understood. In this study, we aimed to investigate the impact of HtrA1 on angiogenic properties of human umbilical vein endothelial cells (HUVECs) as an endothelial model. In particular, the study examined the impact of excessive HtrA1 on capillary tube formation of HUVECs as an *in vitro* angiogenesis model.

## 2. Materials and Methods

### 2.1 In vitro HtrA1 protease activity assay

Full length recombinant human HtrA1 produced in insect cells was from ProteaImmun GmbH (Berlin, Germany), its proteolytic activity was determined by the cleavage of a fluorescence-quenched peptide substrate H2-Opt [Mca-IRRVSYSF(Dnp)KK] [34].

### 2.2 Endothelial tube assay

Human umbilical vein endothelial cells (HUVEC, EA.hy926 from ATCC, Manassas, USA) were maintained in Dulbecco’s Modified Eagle’s Medium (DMEM) and endothelial tube formation was analysed as previously published [35]. Briefly, 96-well cell culture plates were pre-coated with growth factor reduced Matrigel (BD Biosciences, Bedford, MA). HUVECs (12,500 per 100 μl) were plated onto the Matrigel-coated wells, treated with various doses of HtrA1 or vehicle control, and cultured at 37°C for 16 – 20 hours. Cells were then labelled with calcein AM fluorescence dye (BD Biosciences) and assessed through an inverted fluorescent microscope (Olympus, Tokyo, Japan). Images were captured using CellSens software (Olympus); total and mean tube length, branch points and loops were quantified using Image J software (http://rsbweb.nih.gov/ij/; National Institutes of Health, Bethesda, MD).

### 2.3 Statistics

Data are expressed as mean ± SD. Statistical analysis was performed on raw data using one-way ANOVA followed by Tukey’s post hoc test using PRISM (version 5.00, GraphPad Software, San Diego, CA). P<0.05 was taken as significant.

## 3. Results and Discussion

We first confirmed that the recombinant HtrA1 was bioactive as a protease. When HtrA1 was incubated with a fluorescence-quenched peptide substrate at 37°C, a progressive increase in fluorescence signal resulting from substrate cleavage was detected (Figure 1), confirming that HtrA1 was proteolytically activity.

**Figure 1:**
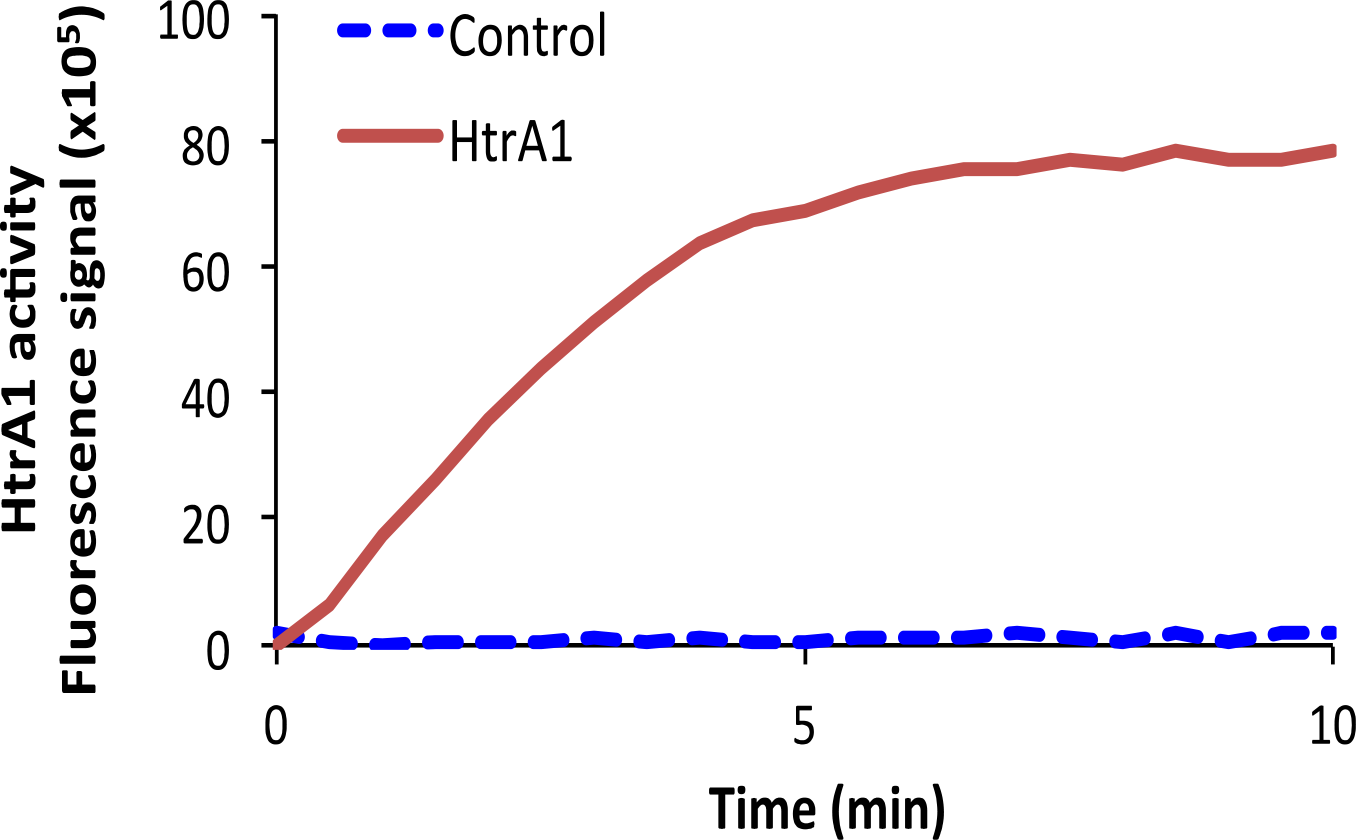
HtrA1 proteolytic activity. Representative real-time progressive curve of substrate cleavage by recombinant human HtrA1.

We then investigated the impact of HtrA1 on HUVEC tube formation. When seeded on growth factor reduced Matrigel, HUVECs formed a regular network of capillary-like tube structure as previously reported (Figure 2A, vehicle control). However, inclusion of HtrA1 induced varying degrees of disruption to HUVEC tube formation. Data of HtrA1 at 1.0, 2.0 and 4.0 ug/ml are shown in Figure 2B-D. The highest concentration of HtrA4 tested was 4.0 ug/ml, as higher than this concentration often caused cells to lift off the plate. When HtrA1 was at 1.0–2.0 μg/ml (Figure 2B-C), HUVECs still formed regular tubes, but they were thinner than controls. However, when HtrA1 concentration was increased to 4.0 ug/ml, HUVECs formed tubes, but they were much smaller and many were incomplete/broken (Figure 2D).

**Figure 2:**
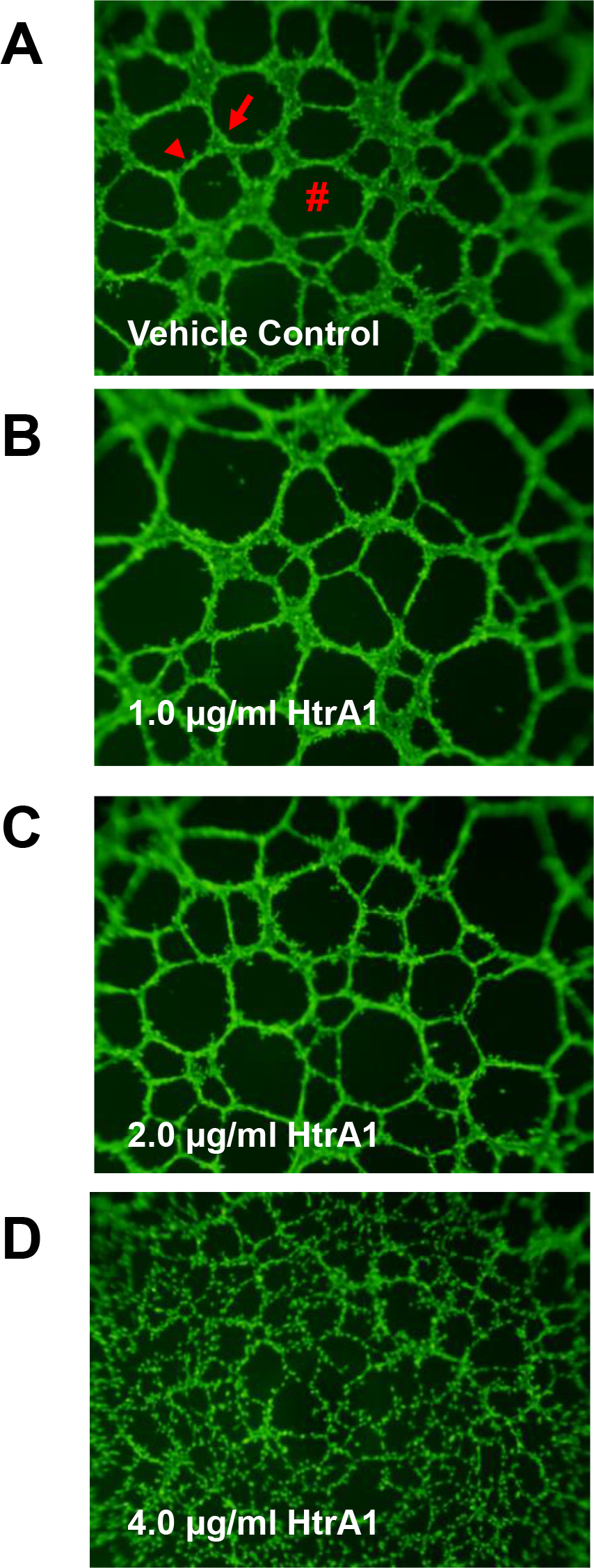
Representative images of HUVEC tube formation in the absence or presence of recombinant HtrA1. (A) Vehicle control. Arrow head, endothelial tube; Arrow, branch point; #, inter-tubular loop. (B-C) HUVECs treated with 1, 2 and 4 μg/ml of recombinant HtrA1 respectively.

To quantify the above characteristics, we measured total tube length (Figure 3A), total branch points (Figure 3B), total number of inter-tubular loops (Figure 3C), and mean tube length (Figure 3D). The highest dose of HtrA1 (4.0 μg/ml) significantly increased the total tube length (Figure 3A), branch points (Figure 3B) and inter-tubular loops (Figure 3C), compared to vehicle control (VC), or the lower doses of HtrA1 (1.0 and 2.0 ug/ml). None of these parameters differed between vehicle control and the lower doses of HtrA1, or between 1.0 and 2.0 ug/ml HtrA1 (Figure 3A-C).

**Figure 2:**
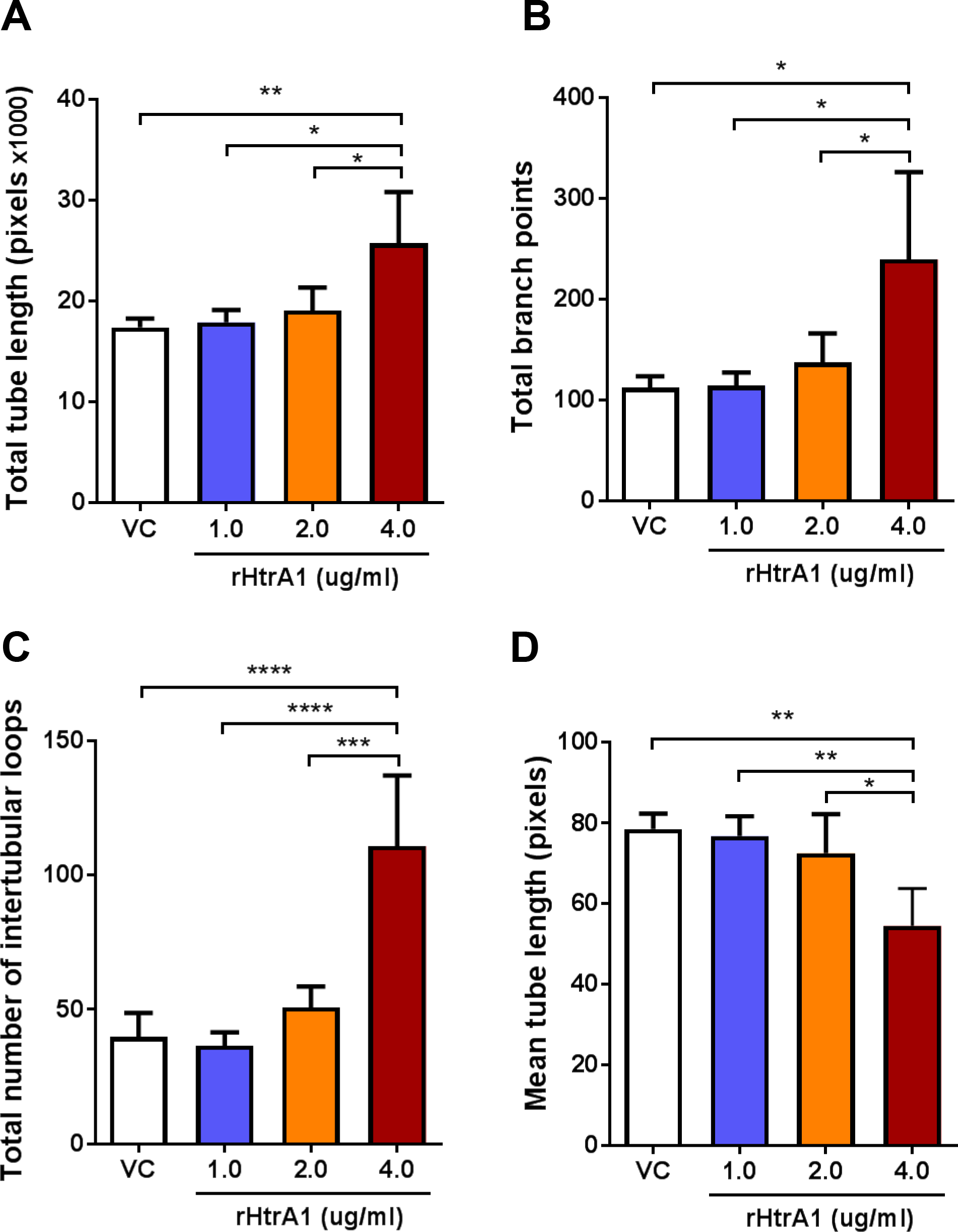
Quantification of the impact of recombinant HtrA1 on HUVEC tube formation. (A) Total tube length, (B) Total branch points, (C) Total inter-tubular loops, and (D) Mean tube length. The data are the average of four independent experiments ± SD. VC, vehicle control. *P<0.05, **P<0.005, ***P<0.0005, ****P<0.0001.

In contrast, the mean length per tube was reduced gradually with increasing concentrations of HtrA1 (Figure 3D). The cells treated with the highest dose of HtrA1 tested (4.0 ug/ml) displayed the shortest mean tube length, which was significantly lower than that of vehicle control (VC), or the lower doses of HtrA1 (1.0 and 2.0 ug/ml). The mean tube length again did not differ between vehicle control and the lower doses of HtrA1, or between 1.0 and 2.0 ug/ml HtrA1 (Figure 3D).

These data indicate that high concentrations of HtrA1 promoted HUVEC tube branching, increased total tube length and total number of inter-tubular loops (Figures 2 and 3). However, the tubes formed were much smaller in size, and importantly they appeared to be incomplete/broken (Figure 2D), suggesting that high concentrations of HtrA1 impaired the completion process of tube formation.

Our studies thus demonstrated that high concentrations of HtrA1 disturbed the angiogenic properties of HUVECs. Although it was an *in vitro* model, these data suggest that elevated HtrA1 observed in both wet AMD and early-onset PE may directly contribute to endothelial dysfunction in these diseases. Future studies are thus warranted to investigate the impact the HtrA1 on microvascular endothelial cells directly involved in the pathogenesis of AMD and PE.

It will also be important to understand the mechanisms of HtrA1 action in inducing endothelial dysfunction. To date, it has been reported that HtrA1 over-expression attenuates cell migration by disrupting microtubule organization, whereas HtrA1 downregulation promotes cell motility [36]. Also, HtrA1 has been suggested to play a critical role in angiogenesis via TGF-β signalling [37]. Recently, HtrA1 was reported to processes extracellular matrix proteins such as thrombospondin 1 in inducing AMD [38]. It is likely that these HtrA1 actions contribute to the disruption of endothelial tube formation, future studies are warranted to investigate the molecular mechanisms of HtrA1 action in inducing endothelial dysfunction, which will provide important insights into the understanding of how elevated levels of HtrA1 contribute to AMD and early-onset PE.

## Acknowledgements

The work was supported by the National Health and Medical Research Council of Australia (Fellowship#1041835 and project grant #1108365 to GN), the Bill and Melinda Gates Foundation, and the Victorian Government Operational Infrastructure Support Program.

